# Matrix-Assisted Laser Desorption-Ionization Time-of-Flight mass spectrometry using a custom-made database, biomarker assignment, or mathematical classifiers does not differentiate *Shigella spp.* and *Escherichia coli*

**DOI:** 10.1101/714295

**Authors:** Maaike J.C. van den Beld, John W.A. Rossen, Arie Evers, A.M.D. (Mirjam) Kooistra-Smid, Frans A.G. Reubsaet

## Abstract

**Purpose:** *Shigella spp*. and *E. coli* are closely related and cannot be distinguished using Matrix-Assisted Laser Desorption-Ionization Time-of-Flight mass spectrometry (MALDI-TOF MS) with commercially available databases. Here, three alternative approaches using MALDI-TOF MS to identify and distinguish *Shigella spp*., *E. coli*, and its pathotype EIEC were explored.

**Methods:** A custom-made database was developed, biomarkers were assigned, and classification models using machine learning were designed and evaluated using spectra of 456 *Shigella spp*., 42 *E. coli*, and 61 EIEC isolates, obtained by the direct smear method and the ethanol-formic acid extraction method. The isolates were identified using ipaH PCR and phenotypic and serological typing.

**Results:** Identification with a custom-made database resulted in >94% *Shigella* identified at the genus level and >91% *S. sonnei* and *S. flexneri* at the species level, but distinction of *S. dysenteriae, S. boydii*, and *E. coli* was poor. Moreover, 10-15% of duplicates rendered discrepant results. With biomarker assignment, 98% *S. sonnei* isolates were correctly identified, although the *S. sonnei* biomarkers were not specific as other species, for instance *S. boydii* and *E. coli* were also identified as *S. sonnei*. Discriminating markers for *S. dysenteriae, S. boydii*, and *E. coli* were not assigned at all. Classifiers identified *Shigella* in 96% of isolates correctly, but most *E. coli* isolates were also assigned to *Shigella*.

**Conclusion:** None of the proposed alternative approaches is suitable for use in clinical diagnostics for the identification of *Shigella spp*., *E. coli*, and EIEC, reflecting their relatedness and problematic taxonomical classification. We suggest the use of MALDI-TOF MS for the identification of the *Shigella spp*./*E. coli* complex, but other tests should be used for distinction.

## Introduction

The *E. coli* pathotype entero-invasive *E. coli* (EIEC) is thought to cause the same disease as *Shigella spp*. [1]. This pathotype consists of isolates that possess some of the *E. coli* phenotypic characteristics and have the invasive nature of *Shigella spp*. [2,3]. EIEC harbors the same virulence markers as *Shigella spp*. that are used in molecular diagnostics to detect both *Shigella spp*. and EIEC but are not suitable to distinguish them [4]. *Shigella spp*. and *E. coli* are described to belong to one taxonomic species genetically, but classification in two genera is maintained for practical and taxonomic reasons [2,5]. Therefore, differentiation is challenging and is historically performed using phenotypical tests, serotyping, and the determination of virulence markers using PCR [6,7].

Most clinical laboratories currently use Matrix-Assisted Laser-Desorption Ionization Time-of-Flight Mass Spectrometry (MALDI-TOF MS) to identify bacteria in a routine diagnostic setting. Commercially available databases, as MALDI biotyper® in combination with the MALDI Security-Relevant (SR) library® (Bruker Daltonik GmbH, Bremen, Germany) and VITEK® MS (BioMérieux, Marcy-l’Etoile, France) can distinguish *Shigella spp*. and *E. coli* from other *Enterobacteriaceae*. However, they cannot distinguish between the different *Shigella* species and *E. coli*, including the EIEC pathotype [8].

The development of custom-made databases to identify bacteria using MALDI-TOF MS as an alternative to commercially available databases proved successful for multiple species before [9-11]. Most notably, an earlier study developed a custom-made database to identify and distinguish *Shigella spp*. and *E. coli* specifically. However, EIEC isolates were not included in their database [12]. Using a database approach, comparisons of unknown isolates to spectra in a database comprise the whole spectra for species identification. However, for closer related groups, a more subtle approach can be essential in which variations within the spectra are examined in the presence or absence of specific peaks as biomarkers [13,14]. The biomarker approach was used to type *E. coli* isolates before, with varying success rates [14]. These approaches mainly targeted a selection of isolates representing the pathotype entero-hemorrhagic *E. coli* (EHEC) or the highly virulent ST131 clone [14], although two studies used biomarker typing specifically for *Shigella spp*. and *E. coli*, without EIEC isolates [15,16]. One of those studies identified biomarkers outside the mass range of 2,000-20,000 Da, which is used in routine applications [16], and the other did not specify in which species the biomarkers were present or absent [15]. Besides determining the presence or absence of single biomarkers, patterns of these biomarkers can be investigated and recognized with machine-learning algorithms [17]. These machine-learning-based methods can establish classifiers for identifying groups within species of bacteria [18,19]. Moreover, these classifiers were developed to identify *Shigella spp*. and *E. coli* before, although EIEC isolates were not included [15].

In this study, the ability of MALDI-TOF MS was assessed for the distinction of the four *Shigella* species, EIEC, and non-invasive *E. coli* using alternatives for the commercially available databases. First, a custom-made database, including all *Shigella* species, *E. coli*, and EIEC isolates, was developed and evaluated. Second, biomarkers were assigned and evaluated, and third, classifier models based on machine learning were defined, applied, and evaluated.

## Material and Methods

### Bacterial isolates

A total of 559 isolates consisting of 36 *S. dysenteriae*, 156 *S. flexneri*, 32 *S. boydii*, 232 *S. sonnei*, 61 EIEC, and 42 other *E. coli* of human and animal origin comprising phylogroups A, B1, B2, and D [20] was used (Table 1). All isolates, except the references, were identified using a previously described culture-based identification algorithm [21]. They were divided into a set of training isolates (n=288) and test isolates (n=271), both having similar species and serotype distributions. The training set was used to construct the custom-made database, assign biomarkers, and define and train machine-learning classifier models. The test set was used to test all of these algorithms in duplicate, with both direct smear and ethanol-formic acid extraction application methods.

**Table 1.** Isolates used in this study, divided in training set and test set.

| Species and serotype/O-type | Training set |  | Test set |  |
| --- | --- | --- | --- | --- |
|  | n | Origin | n | Origin |
| *S. dysenteriae* serotype 1 | 2 | CIP 57.28<sup>T</sup>; A1 | 1 | 1 ci<sup>a</sup> |
| *S. dysenteriae* serotype 2 | 5 | A2, 4 ci<sup>a</sup> | 4 | 4 ci<sup>a</sup> |
| *S. dysenteriae* serotype 3 | 5 | AMC-43-G-93; 4 ci<sup>a</sup> | 3 | 3 ci<sup>a</sup> |
| *S. dysenteriae* serotype 4 | 2 | AMC 43-G-86; 1 ci<sup>a</sup> | 0 |  |
| *S. dysenteriae* serotype 5 | 1 | AMC 43-G-84 | 0 |  |
| *S. dysenteriae* serotype 6 | 1 | AMC 43-G-81 | 1 | 1 ci<sup>a</sup> |
| *S. dysenteriae* serotype 7 | 1 | AMC 43-G-76 | 1 | 1 ci<sup>a</sup> |
| *S. dysenteriae* serotype 9 | 2 | A58: 1646;<br>1 ci<sup>a</sup> | 1 | 1 ci<sup>a</sup> |
| *S. dysenteriae* serotype 10 | 1 | A2050-52 | 0 |  |
| *S. dysenteriae* serotype 12 | 2 | 2 ci<sup>a</sup> | 1 | 1 ci<sup>a</sup> |
| *S. dysenteriae* serotype 14 | 1 | NCTC 11867 | 0 |  |
| *S. dysenteriae* serotype 15 | 1 | NCTC 11868 | 0 |  |
| **Total number of *S. dysenteriae*** | **24** |  | **12** |  |
| *S. flexneri* serotype 1a | 3 | B1A; 2 ci<sup>a</sup> | 0 |  |
| *S. flexneri* serotype 1b | 5 | B1B; 4 ci<sup>a</sup> | 5 | 5 ci<sup>a</sup> |
| *S. flexneri* serotype 1c | 4 | 4 ci<sup>a</sup> | 3 | 3 ci<sup>a</sup> |
| *S. flexneri* serotype 2a | 32 | CIP 82.48<sup>T</sup>; B2A; 30 ci<sup>a</sup> | 32 | 32 ci<sup>a</sup> |
| *S. flexneri* serotype 2b | 1 | B2B | 3 | 3 ci<sup>a</sup> |
| *S. flexneri* serotype 3a | 2 | B3A; 1 ci<sup>a</sup> | 14 | 14 ci<sup>a</sup> |
| *S. flexneri* serotype 3b | 2 | B3B; B3C | 3 | 3 ci<sup>a</sup> |
| *S. flexneri* serotype 4a | 1 | B4A | 4 | 4 ci<sup>a</sup> |
| *S. flexneri* serotype 4av | 4 | 5 ci<sup>a</sup> | 0 |  |
| *S. flexneri* serotype 4b | 1 | B4B | 0 |  |
| *S. flexneri* serotype 4c | 3 | 3 ci<sup>a</sup> | 0 |  |
| *S. flexneri* serotype 5b | 1 | B5 | 1 | 1 ci<sup>a</sup> |
| *S. flexneri* serotype 6 | 10 | B6; 9 ci<sup>a</sup> | 10 | 10 ci<sup>a</sup> |
| *S. flexneri* serotype X | 1 |  |  |  |
| *S. flexneri* serotype Y | 2 | 2 ci<sup>a</sup> | 2 | 2 ci<sup>a</sup> |
| *S. flexneri* serotype Yv | 2 | 2 ci<sup>a</sup> | 0 |  |
| *S. flexneri* provisional | 5 | 5 ci<sup>a</sup> | 0 |  |
| **Total number of *S. flexneri*** | **79** |  | **77** |  |
| *S. boydii* serotype 1 | 2 | AMC-43-G-58; 1 ci<sup>a</sup> | 3 | 3 ci<sup>a</sup> |
| *S. boydii* serotype 2 | 3 | CIP 82.50<sup>T</sup>; P288; 1 ci<sup>a</sup> | 4 | 4 ci<sup>a</sup> |
| *S. boydii* serotype 3 | 1 | D1 | 0 |  |
| *S. boydii* serotype 4 | 2 | AMC-43-G-63; 1 ci<sup>a</sup> | 2 | 2 ci<sup>a</sup> |
| *S. boydii* serotype 5 | 2 | P143; 1 ci<sup>a</sup> | 0 |  |
| *S. boydii* serotype 6 | 1 | CDC 9771 (D19) | 0 |  |
| *S. boydii* serotype 7 | 1 | AMC 4006 (Lavington) | 0 |  |
| *S. boydii* serotype 8 | 0 |  | 1 | 1 ci<sup>a</sup> |
| *S. boydii* serotype 9 | 1 | 1296/7 | 0 |  |
| *S. boydii* serotype 10 | 1 | 430 | 1 | 1 ci<sup>a</sup> |
| *S. boydii* serotype 11 | 1 | 34 | 0 |  |
| *S. boydii* serotype 12 | 0 |  | 1 | 1 ci<sup>a</sup> |
| *S. boydii* serotype 13 | 0 |  | 1 | 1 ci<sup>a</sup> |
| *S. boydii* serotype 14 | 0 |  | 1 | 1 ci<sup>a</sup> |
| *S. boydii* serotype 15 | 1 | CDC C-703 | 0 |  |
| *S. boydii* serotype 18 | 1 | 1 ci<sup>a</sup> | 1 | 1 ci<sup>a</sup> |
| **Total number of *S. boydii*** | **17** |  | **15** |  |
| *S. sonnei* | 117 | CIP 82.49<sup>T</sup>;<br>116 ci<sup>a</sup> | 115 | 115 ci<sup>a</sup> |
| EIEC | 30 | DSM 9027; DSM 9028; CCUG 11335;<br>CCUG 38080; CCUG 38092; CCUG<br>38093; EW227; 1624-56; 1184-68;<br>145/46; L119B-10; 19 ci<sup>a</sup> | 31 | 31 ci<sup>a</sup> |
| Other *E. coli* pathotypes<br>(human) | 11 | 7 STEC ci<sup>a</sup>, 4 EPEC ci<sup>a</sup> | 11 | 8 STEC ci<sup>a</sup>; 3 EPEC ci<sup>a</sup> |
| Other *E. coli* pathotypes<br>(animal)<sup>b</sup> | 10 | 5 mussel, 3 pigeon, 2 turkey | 10 | 4 mussel, 3 pigeon, 2 turkey, 1<br>oyster |

<sup>a</sup>ci = clinical isolate. <sup>b</sup>isolated from animals, all other numbers are reference isolates.

### MALDI-TOF MS preparation of isolates

All isolates were grown overnight on Columbia Sheep Agar (CSA, Biotrading, Mijdrecht, the Netherlands) at 37°C and were subsequently subjected to the direct smear method and the ethanol-formic acid extraction with silica beads as previously described [22]. Colonies or 1 µl extract were applied onto a polished steel plate, air-dried, and overlaid with 1 µl α-Cyano-4-hydroxycinnamic acid in 50% acetonitrile-2.5% trifluoroacetic acid (HCCA matrix). The samples were analyzed using a Bruker Microflex LT (Bruker Daltonik GmbH, Bremen, Germany) in a linear and positive mode, with 30-40% laser power and within a mass-range of 2,000-20,000 Da.

The isolates from the training set were used to produce Main Spectrum Profiles (MSPs) according to the manufacturer’s instructions, using FlexControl V3.4 (Bruker Daltonik). The isolates from the training set were analyzed using Maldi Biotyper 3.0 Real-Time Classification (RTC, Bruker Daltonik).

### Database development

The MSPs produced from 288 isolates in the training set were used to build a custom-made database with Maldi Biotyper OC V3.1.66 (Bruker Daltonik). In addition, a dendrogram to assess the relatedness of these MSPs was inferred using default settings. The isolates in the test set were identified using this custom-made database. Additionally, the test isolates were also identified using the commercially available Bruker MALDI Biotyper database (V8.0.0.0) and the Bruker Security-Relevant Library (V1.0.0.0) and using a combination of the commercial and custom-made databases. Quality of the results was indicated by a log-score, calculated by Maldi Biotyper 3.0 RTC: a log-score of 2.000-2.300 corresponds to “secure genus identification, probable species identification,” and a log-score of >2.300 corresponds to a “highly probable species identification.” Both duplicate spots were analyzed, the highest log-score of at least 2.000 was considered as the definitive MALDI-TOF MS identification, as is done in a routine workflow. If an isolate had a log-score < 2.000 caused by a poor spectrum, it was disregarded from further analysis. Isolates were then assigned to different discrimination levels “genus,” “pathotype,” “group,” and “species,” as displayed in Figure 1.

**Fig. 1.**
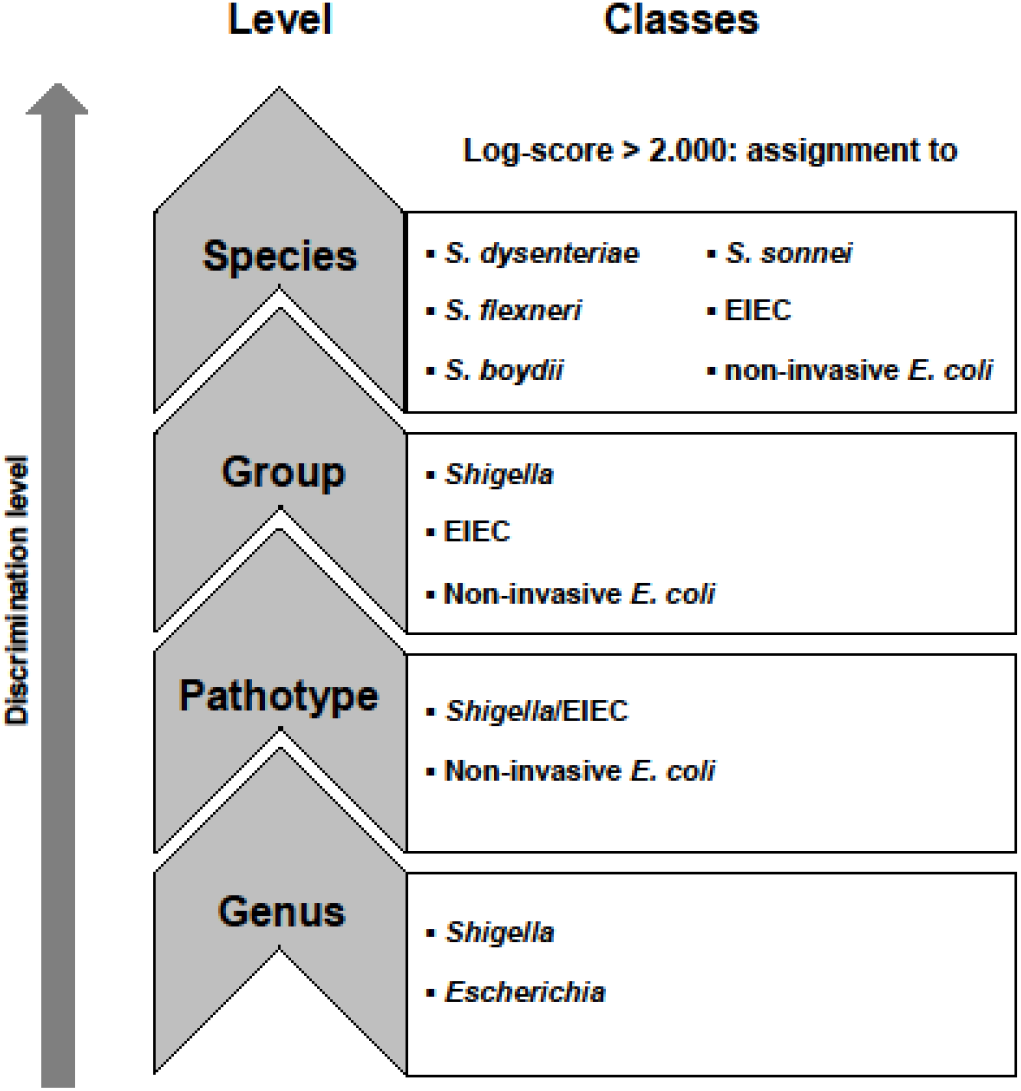
The classes in the different discrimination levels to which isolates were assigned.

For accurate identification, only matches with database MSPs from the same species within a log-score range of 2.000-2.300 or >2.300 should be expected in one spot. Therefore, the ten MSPs from the database that produced the highest scores within a log-score of 2.000-2.300 or >2.300 per spot were determined. For each species identified with the culture-dependent identification algorithm, the median number of species resulting from MALDI-TOF MS and their quartile ranges per spot that had a log-score of 2.000-2.300, or >2.300 were calculated and visualized using SPSS 24.0.0.1 (IBM, New York, USA).

### Biomarker assignment and principal component analysis

Spectra files from MSPs of 288 isolates in the training set were exported as mzXML files using Compassxport CXP3.0.5. (Bruker Daltonik) or exported via a batch process in Flexanalysis (Bruker Daltonik). A new database was created in Bionumerics v7.6.3 (Applied Maths NV, http://www.applied-maths.com/) according to the manufacturers’ instructions. All raw spectra were imported into the Bionumerics database with x-axis trimming to a minimum of 2000 m/z. Baseline subtraction, noise computing, smoothing, baseline detection, and peak detection were performed with default settings. Spectra summarizing, peak matching, and peak assignment were performed according to instructions from Bionumerics [23]. In short, all raw spectra were summarized into isolate spectra. Peak matching was performed on isolate spectra using a constant tolerance of 1.9, a linear tolerance of 550, and a peak detection rate of 10%. Binary peak matching tables were exported to summarize the presence of peak classes on all discrimination levels, as depicted in Figure 1. For the levels genus, pathotype and groups, decision diagrams were produced (Supplementary file 1a, 1b, and 1c). The spectra files of isolates from the test set were imported and preprocessed in Bionumerics, using the same methods and settings as for the spectra from the isolates in the training set. Peak matching of test isolates was performed using the option “existing peak classes only” to compare the presence of peaks in the test isolates with peaks in the isolates from the training set. Decision diagrams (Supplementary File 1) and the presence or absence of peak masses as depicted in Table 2 were applied to assign unknown isolates from the test set according to the different levels, as shown in Figure 1.

**Table 2.**
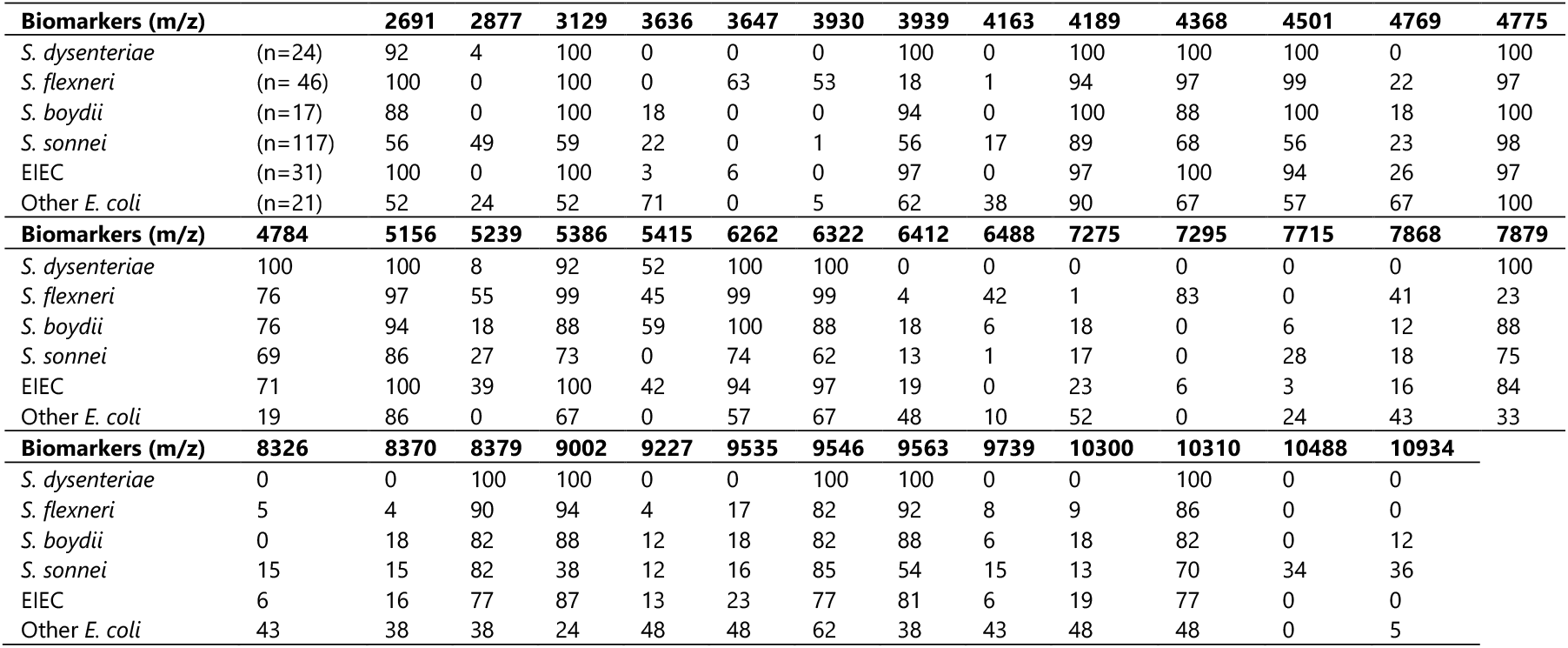
Discrimination scheme of biomarkers, percentage of isolates in the training set with presence of specific biomarkers.

By assigning biomarkers, only the presence and absence of peaks were investigated. To assess quantitative peak data such as peak intensity and peak area, a principal component analysis (PCA) was performed on all isolates in the training set to visualize the position of isolates in three dimensions.

### Presence of biomarkers identified in previous studies

All isolates in the training set and the test set were examined for the unique masses (± 500 ppm) found in biomarker assignment to *Shigella spp*. and *E. coli* in previous studies [16,15]. Additionally, because peaks in our study were assigned at m/z values instead of masses only and masses could be potentially charged with two electrons, this is corrected by examining the previously published masses divided by two (±500 ppm) [15,16].

### Classifier models based on machine learning

Peak data of the summarized isolate spectra of the 288 isolates in the training set was used to define and train machine learning-based classifiers using Bionumerics v7.6.3 according to the manufacturers’ instructions. In short, peak matching with a constant tolerance of 1.9 and a linear tolerance of 550 was performed on isolate spectra on the different levels: genus, pathotype, group, and species. Classifiers were created at all levels using character values. Support Vector Machine (linear) learning was used as a scoring method in which p-values were used to rank. The classifiers were trained and cross-validated to check their performance for identification. Subsequently, the classifier models were used to classify the unknown isolates in the test set at the different discrimination levels to evaluate their performance.

## Results

### Database development

All MSPs of 288 training isolates were added to a custom-made database. The relatedness of these MSPs is shown in a dendrogram (Figure 2). The Maldi Biotyper OC software recognized three large MSPs clusters that are not species-specific within this custom database. This did not change if clusters were assigned manually with a lower distance level at 50-100 relative units, indicating that similarity in spectrum profiles is distributed over the species level (Figure 2).

**Fig. 2.**
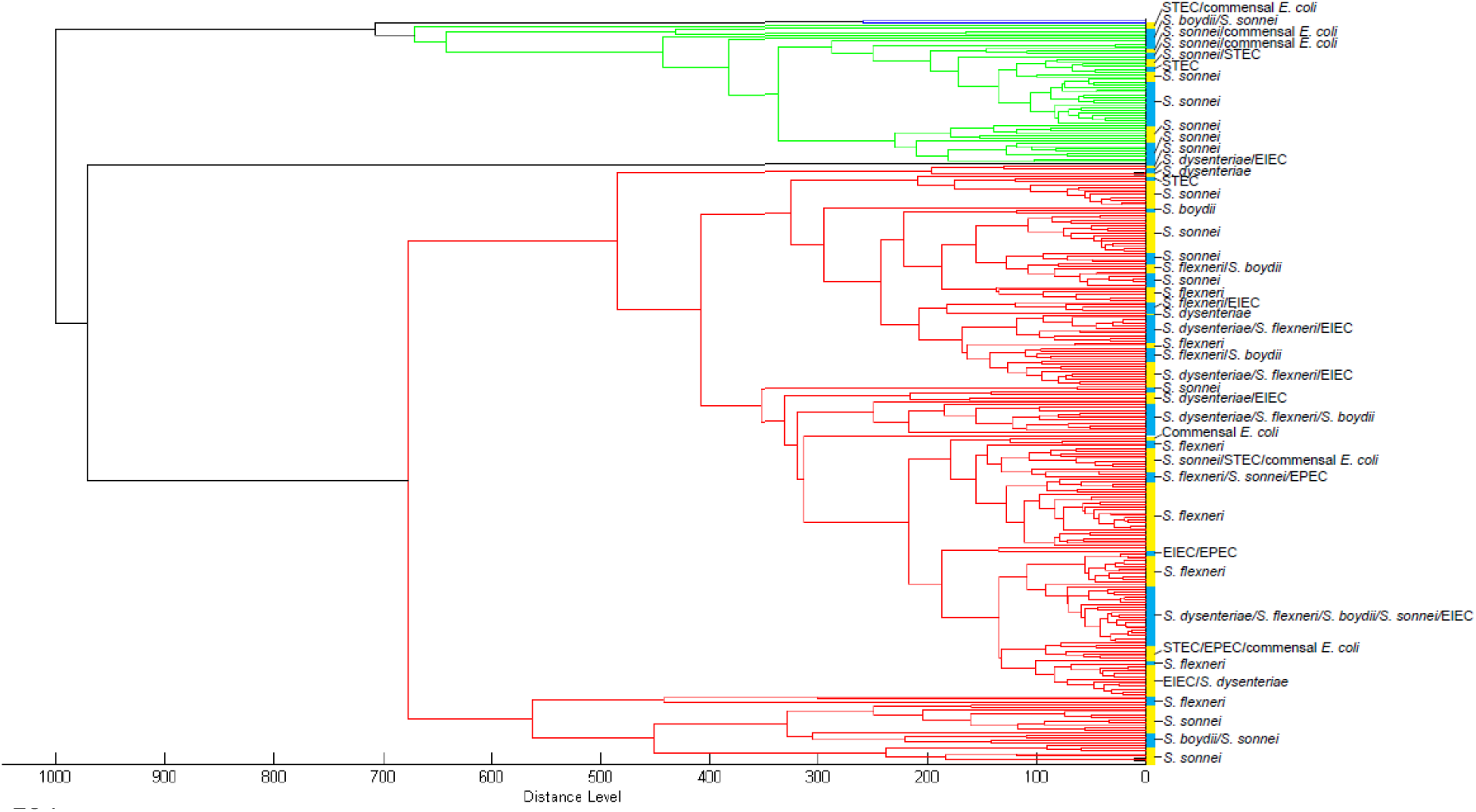
Dendrogram of MSPs of training isolates. *Blue = cluster 1; green = cluster 2; red = cluster 3. Yellow/blue vertical band = manual cluster distinction at distance level 50-100 relative units with species designation using the culture-based identification algorithm*.

Additionally, the duplicate spots of test isolates using either the direct smear or extraction method resulted in a different species designation in 10-15% of the samples. Furthermore, with an accurate distinction of species, one would not expect assignment to multiple species above the threshold of log-score 2.000. However, with both application methods, most isolates were assigned to several species with a log-score of 2.000-2.300 or >2.300 per spot (Figure 3).

**Fig. 3.**
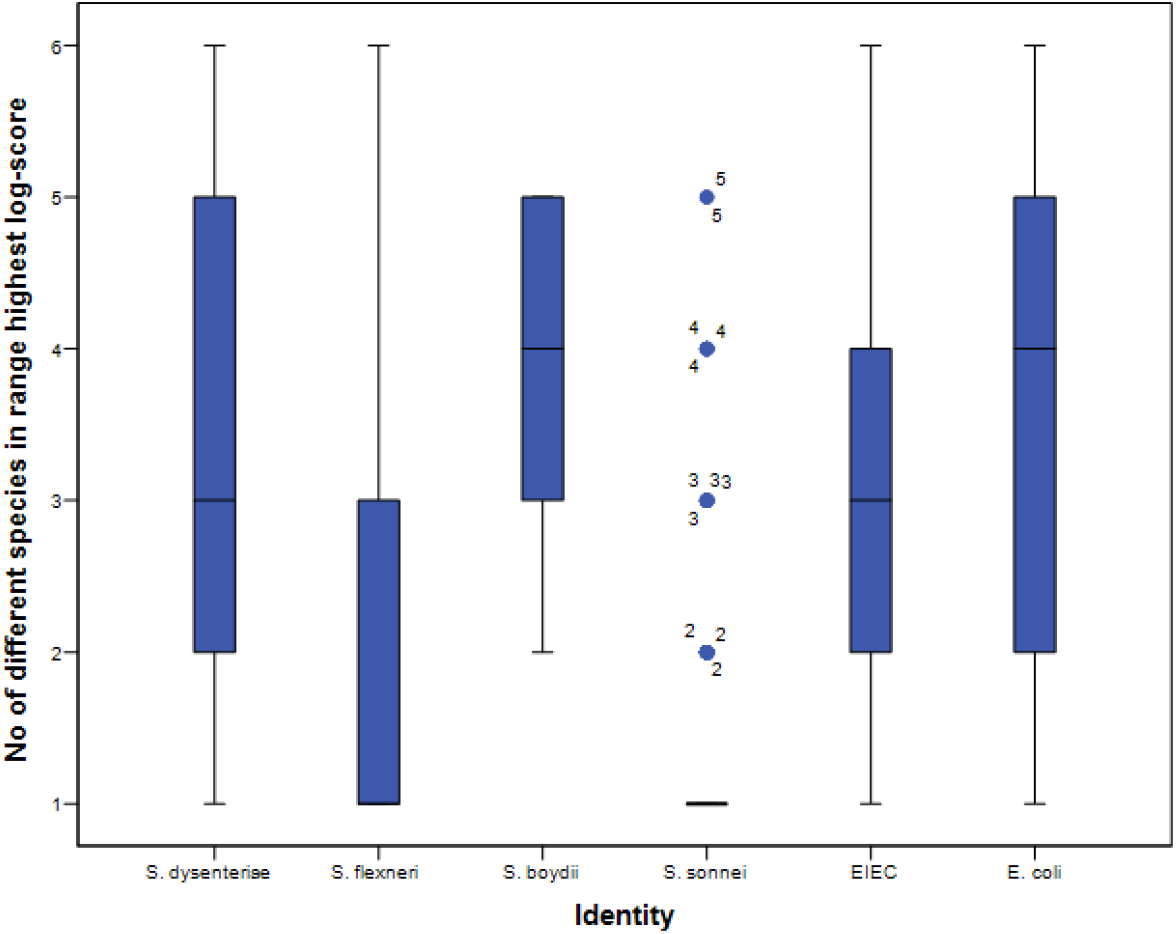
Number of different species in the first 10 matches per spot with the direct smear method. *Identity (x-axis) was assigned using the culture-based identification algorithm. Black horizontal bars represent the median number of species, the 25%-75% interquartile ranges are indicated by the blue vertical bars, and 5%-95% intervals by the black vertical lines. Outliers are indicated with blue dots*.

One isolate from the test set (*S. boydii* serotype 13) had a low-quality spectrum (log score 1.574-1.930), and one isolate *(S. dysenteriae* serotype 1) has initially been incorrectly stored, as this isolate was identified as *Corynebacterium diphtheria*e using the Bruker databases. Both these isolates were ignored in further analyses. All other isolates had log-scores higher than 2.000, and percentages of MALDI-TOF MS identification concordant with the original identification on all discrimination levels were as displayed in Table 3. With the Bruker databases only, percentages of correctly identified *Shigella spp*. on all discrimination levels are low, ranging from 6% to 45%correct designations, both for the direct smear method and the extraction method (Table 3). In contrast, 90%-100% of *E. coli* isolates were correctly identified. When identification was based on the custom-made database with or without the Bruker databases, the percentage of correctly identified *E. coli* isolates decreased to a range of 29%-71%, while *Shigella spp*. were correctly identified, ranging from 94% to99% of cases on the genus, pathotype and group levels. In addition, 91%-97% of *S. flexneri* and *S. sonnei* were correctly identified at the species level, in contrast to *S. dysenteriae* and *S. boydii*, for which the percentages of correct identification were low (Table 3).

**Table 3.**
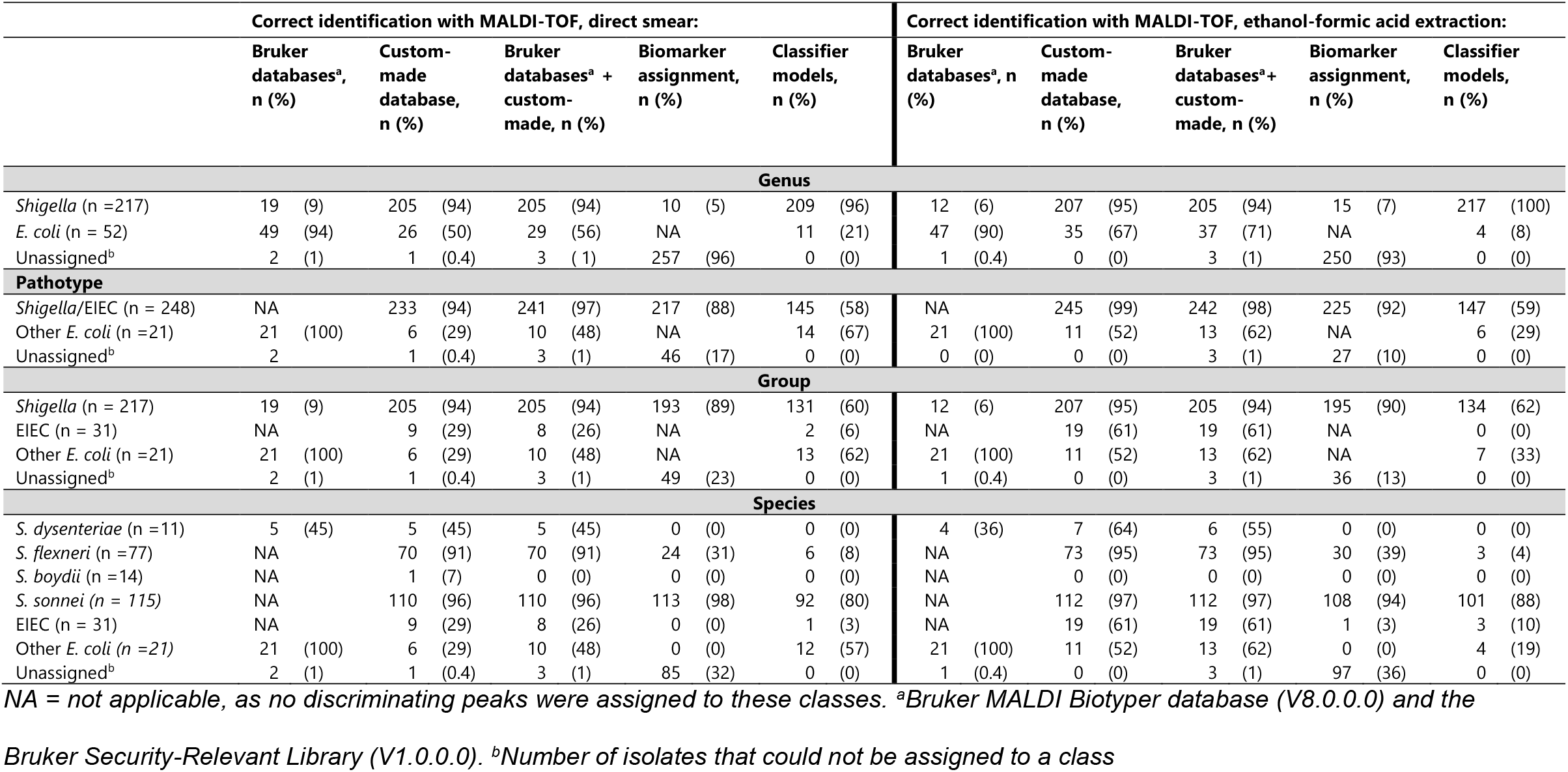
Correct identification results of isolates from test set.

### Biomarker assignment and principal component analysis

The decision diagrams based on biomarkers assigned to the isolates in the training set were used for the identification of unknown isolates in the test set. Distinctive peaks on the species levels were summarized in Table 2. High percentages for correct identification of *S. sonnei* isolates were achieved at the species level using both the direct smear as the extraction method. However, the biomarkers are not specific for *S. sonnei* as other species also contain them. For other species, the identified biomarkers correctly identified isolates below 38%. Specific biomarkers were not detected for all the classes at the different discrimination levels, as depicted in Figure 1. Consequently, it was not possible to identify *S. dysenteriae, S. boydii*, and *E. coli* isolates at all because of the absence of discriminating peaks for these species (Table 3).

In the PCA of the detected peaks in the isolates of the training set, one large cluster was formed, with a few outliers at both ends (Figure 4). If the isolates were colored according to their identity based on the culture-based identification method, separate groups of isolates were seen in none of the discrimination levels (Figure 4a-d).

**Fig. 4.**
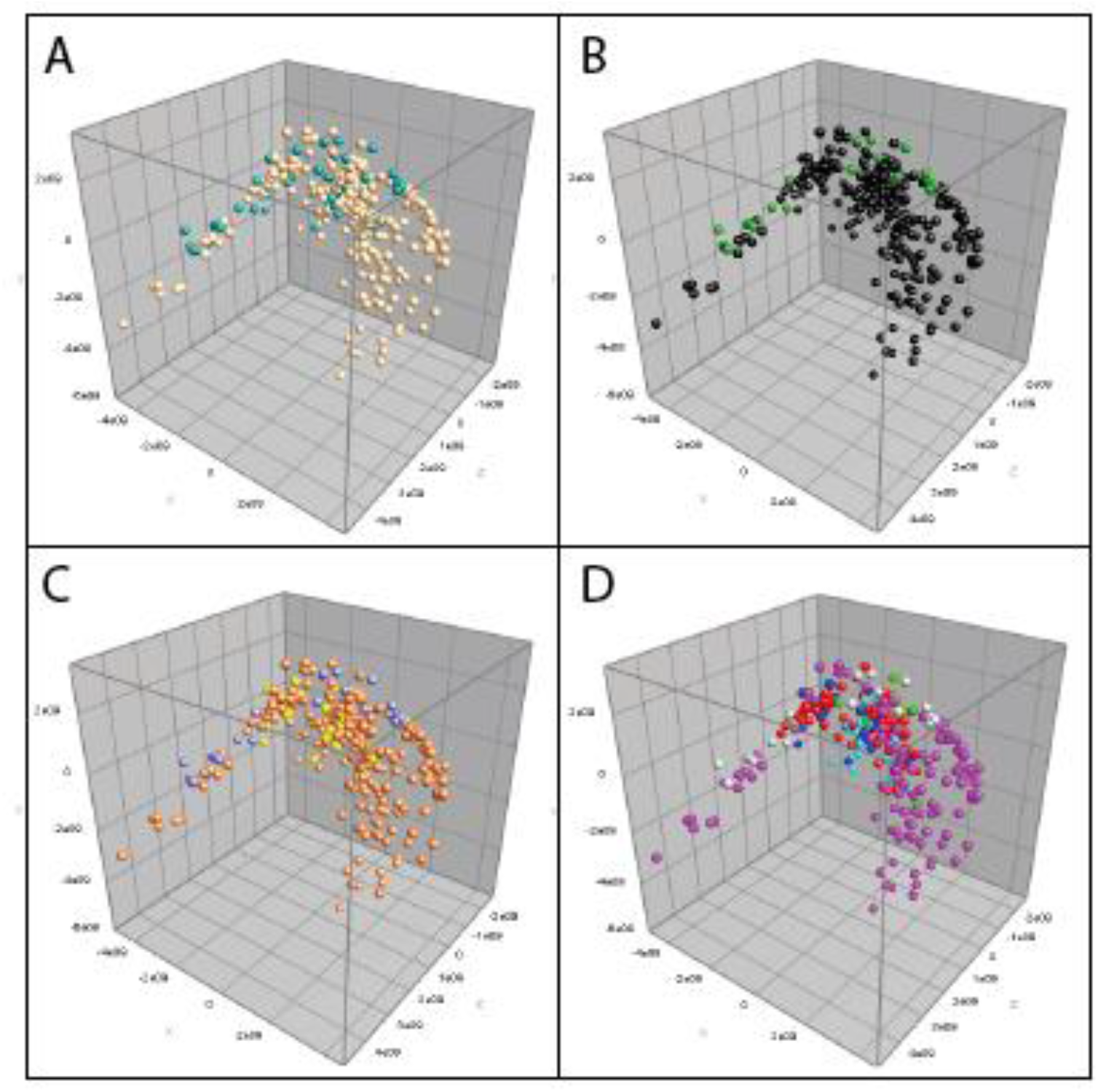
PCA of isolates in the training set. *a. Colored at genus level: beige = Shigella; teal = Escherichia. b. Colored at pathotype level: black = Shigella/EIEC; green = E. coli (other than EIEC). c. Colored at group level: orange = Shigella spp*.; *yellow = EIEC; purple = E. coli (other than EIEC). d. Colored at species level: light blue = S. dysenteriae; red = S. flexneri; green = S. boydii; pink = S. sonnei; blue = EIEC; light grey = Other E. coli*.

### Presence of biomarkers identified in previous studies

The specific biomarkers for *S. flexneri, S. sonnei*, and *E. coli* assigned by Everley *et al*. [16] were not present in any of the 559 isolates in this study when using an error limit of ± 500 ppm. If they were corrected for a charge with 2 electrons, they were also not present. A few biomarkers for *Shigella spp*. and *E. coli* described by Khot and Fisher [15] were present within a range of 500 ppm in isolates used in this study, i.e., 4163 Da, 7157 Da, 8326 Da, and 9227 Da, and corrected for a charge of 2 electrons, 5096 Da and 5752 Da.

### Classifier models based on machine learning

Using the internal cross-validation of the classifiers at all discrimination levels, all but one class had an accuracy of more than 87.5%. The only class with a lower accuracy (77%) was the class “*Escherichia*” at the genus discrimination level.

When using machine learning-based classifiers for identification, 96% of *Shigella spp*. isolates and 21% of the *E. coli* isolates from the test set were correctly identified at the genus level, using the direct smear application method and respectively 100% and 8% using the ethanol-formic acid extraction method (Table 3). Correct identification percentages for the pathotype, group, and species level were displayed in Table 3. Although more than 80% of *S. sonnei* isolates were correctly identified with the species classifier, specificity is low, as more than 70% of *S. flexneri* isolates were also identified as *S. sonnei*.

## Discussion

Current commercially available MALDI-TOF MS databases cannot distinguish between *Shigella spp*. and *E. coli*. Therefore, three different alternatives were explored in this study. A custom-made database was developed, biomarkers were identified, and classification models using machine learning were designed.

Compared to a previous study, our custom-made database assigned fewer *E. coli* isolates correctly [12]. This indicates that the inclusion of EIEC isolates in the custom-made database and the test set complicates the identification. Half of the EIEC isolates were assigned to one of the *Shigella* species, thereby decreasing the percentage of correctly identified *E. coli*. The poor performance of identifying *E. coli* with our custom-made database can result from an overrepresentation of *S. flexneri* and *S. sonnei*. A second custom-made database was developed to investigate this hypothesis, based on 17 isolates of each species, representing the diversity in serotypes. This database did not perform better or worse than the custom-made database that contained 288 MSPs (Supplementary File 2), indicating that a more evenly distribution of species in the database does not improve the identification of *E. coli*. Although percentages of correct species assignments to *S. flexneri* and *S. sonnei* were high, other species were falsely assigned to them, both in our study as in a previous study [12]. In the latter study, correct species identification was based on the majority rule that three out of four spots should indicate the same species. Besides the fact that the interpretation of four spots per isolate is not feasible in clinical diagnostics, this indicates that the assignment of species is based on probabilities rather than actual variations in spectra. Our study confirms this phenomenon because multiple species identifications within the same log-score range were made per spot. Moreover, 10-15% of duplicate spots resulted in different species assignments using the commercially available databases and the custom-made database. Additionally, in the dendrogram that was inferred from the MSPs in the custom-made database, the same species were not clustering together, indicating that the resulting database would not be capable of identifying the isolates from the test set correctly.

Another alternative approach for the use of commercially available databases is the detection of discriminating biomarkers. However, in our study, many isolates had an inconclusive identification, as specific biomarkers were not detected for most classes. Although more than 90% of *S. sonnei* isolates were identified at the species level, other species as *S. boydii* and *E. coli* are also frequently falsely identified as *S. sonnei*. Moreover, when also analyzing peak intensity and peak area rather than just peak presence, the PCA showed that *Shigella spp*. and *E. coli* did not represent separated groups based on their biomarkers. In contrast, one large cluster with a few outliers was formed, demonstrating their genetic similarity. Furthermore, the absence of 85% of the masses assigned as biomarkers in a former study in our isolates [15] indicates that the detected biomarkers vary amongst isolate sets tested and that a stable variation per species is not observed. Consequently, we anticipate that the assignment of biomarkers based on yet another additional set of isolates will lead to even more diversity in biomarkers, demonstrating their unsuitability for distinct identification of *Shigella spp*., *E. coli*, and EIEC.

The use of classifier models based on machine learning resulted in comparable percentages of correctly identified *Shigella* on the genus level, i.e., ≥ 94%, as reported in other studies [15]. In our classifier model designed on the pathotype level, EIEC isolates were not incorporated in the class *E. coli*, and correct identification was 67%, comparable to a previous study [15]. Nonetheless, the other remaining *E. coli* isolates were falsely classified as *Shigella* both with our classifiers and with previously published ones [15], decreasing the specificity for the identification of *Shigella*. At the group and species level, classifiers performed even less, and most species could not be identified at all. The poor performance of the classifier models may be caused by an overrepresentation of *S. flexneri* and *S. sonnei*, as discussed for the custom-made database in our study. Therefore, the selection of 17 isolates of each species was used again, and alternative classifiers were designed. These classifiers did not perform better or worse than the classifiers designed using all 288 isolates in the training set, indicating that an absence of an evenly distribution of species was not the cause for poor identification with classifiers (Supplementary File 2).

Compared to previous studies, we used a substantially more extensive set of isolates and included the *E. coli* pathotype EIEC. Another strength of our study was that multiple alternative approaches for identifying *Shigella spp*. and *E. coli* using MALDI-TOF MS were explored. Although *S. sonnei* and *S. flexneri* isolates were overrepresented in both the training set and the test set, this distribution represents high-resource settings.

In conclusion, none of our explored alternative approaches for identifying *Shigella spp*., *E. coli*, and EIEC with MALDI-TOF MS was suitable to use in clinical diagnostics as all rendered a poor distinction based on spectra or biomarkers. This poor discrimination merely reflects the problematic taxonomical classification of *Shigella spp*. and *E. coli* into two different genera and does not reflect MALDI-TOF MS’s performance as an identification technique in general. Therefore, we propose an identification algorithm in which MALDI-TOF MS is used to identify and differentiate *Shigella/E. coli* as a group from other *Enterobacteriaceae*, followed by tests other than MALDI-TOF MS to distinguish between the different *Shigella species, E. coli*, and specific *E. coli* pathotypes, including EIEC.

## Author contributions

MB, JR, MK, and FR conceived and designed the project. MB and AE performed the analyses. MB, JR, MK, and FR interpreted results. MB wrote the manuscript. All authors read, reviewed, and approved the final manuscript.

## Data availability statement

The datasets generated during and/or analyzed during the current study are available from the corresponding author on reasonable request.

## Supplementary Material

Supplementary File_1.pdf Decision diagrams of assigned biomarkers Supplementary File_2.pdf Comparison of performance of a custom-made database and classifiers based on all 288 isolates or based on an evenly distribution of 17 isolates for each species

