## Supplemental material for "Matrix-Assisted Laser Desorption-Ionization Time-of-Flight mass spectrometry using a custom-made database, biomarker assignment, or mathematical classifiers does not differentiate *Shigella spp.* and *Escherichia coli*"

**Supplementary File 1** Decision diagrams of assigned biomarkers

a.Biomarkers at genus level. b. Biomarkers at pathotype level. c. Biomarkers at group level

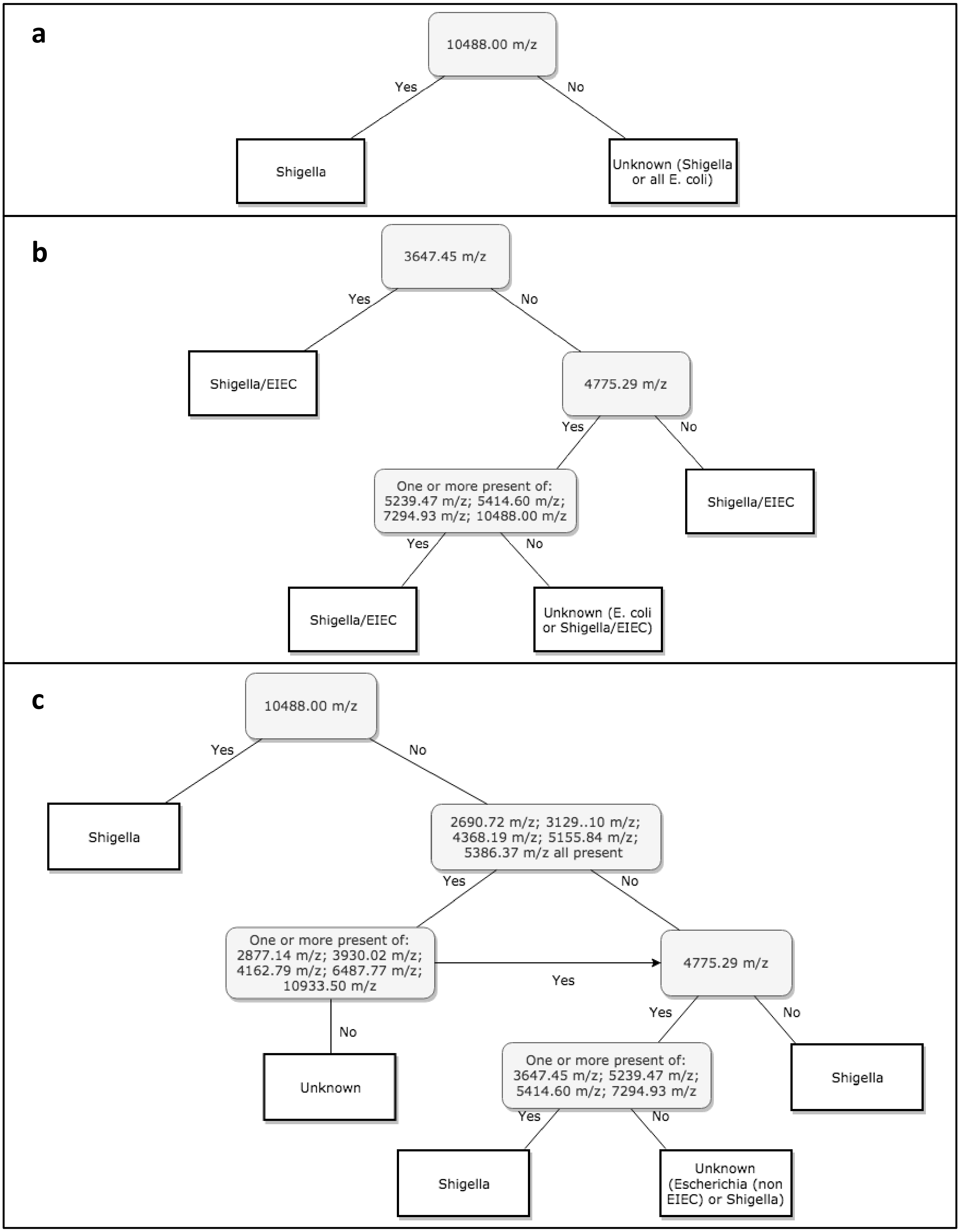

**Supplementary File 2** Comparison of performance of a custom-made database and classifiers based on all 288 isolates or based on an evenly distribution of 17 isolates for each species (6*17)

| | Correct identification with MALDI-TOF, direct smear: | | | | Correct identification with MALDI-TOF, ethanol-formic acid extraction: | | | |
|---|---|---|---|---|---|---|---|---|
| | **Custom-made database, training set 288 isolates n (%)** | **Custom-made database, training set 6*17 isolates n (%)** | **Classifier models, training set 288 isolates n (%)** | **Classifier models, training set 6*17 isolates n (%)** | **Custom-made database, training set 288 isolates n (%)** | **Custom-made database, training set 6*17 isolates n (%)** | **Classifier models, training set 288 isolates n (%)** | **Classifier models, training set 6*17 isolates n (%)** |
| **Genus** | | | | | | | | |
| ***Shigella* (n =217)** | 205 (94) | 196 (90) | 209 (96) | 157 (72) | 207 (95) | 201 (93) | 217 (100) | 191 (88) |
| ***E. coli* (n = 52)** | 26 (50) | 28 (54) | 11 (21) | 11 (21) | 35 (67) | 27 (52) | 4 (8) | 9 (17) |
| **Unassigned** | 1 (0.4) | 1 (0.4) | 0 (0) | 0 (0) | 0 (0) | 2 (1) | 0 (0) | 0 (0) |
| **Pathotype** | | | | | | | | |
| ***Shigella*/EIEC (n = 248)** | 233 (94) | 222 (90) | 145 (58) | 157 (63) | 245 (99) | 241 (97) | 147 (59) | 242 (96) |
| **Other *E. coli* (n =21)** | 6 (29) | 10 (48) | 14 (67) | 1 (5) | 11 (52) | 9 (43) | 6 (29) | 1 (5) |
| **Unassigned** | 1 (0.4) | 1 (0.4) | 0 (0) | 0 (0) | 0 (0) | 2 (1) | 0 (0) | 0 (0) |
| **Group** | | | | | | | | |
| ***Shigella* (n = 217)** | 205 (94) | 196 (90) | 131 (60) | 142 (65) | 207 (95) | 201 (93) | 134 (62) | 208 (96) |
| **EIEC (n = 31)** | 9 (29) | 5 (16) | 2 (6) | 1 (3) | 19 (61) | 11 (35) | 0 (0) | 1 (3) |
| **Other *E. coli* (n =21)** | 6 (29) | 10 (48) | 13 (62) | 1 (5) | 11 (52) | 9 (43) | 7 (33) | 1 (5) |
| **Unassigned** | 1 (0.4) | 1 (0.4 | 0 (0) | 0 (0) | 0 (0) | 2 (1) | 0 (0) | 0 (0) |
| **Species** | | | | | | | | |
| ***S. dysenteriae* (n =11)** | 5 (45) | 5 (45) | 0 (0) | 0 (0) | 7 (64) | 7 (64) | 0 (0) | 1 (9) |
| ***S. flexneri* (n =77)** | 70 (91) | 59 (77) | 6 (8) | 0 (0) | 73 (95) | 64 (83) | 3 (4) | 0 (0) |
| ***S. boydii* (n =14)** | 1 (7) | 4 (29) | 0 (0) | 0 (0) | 0 (0) | 3 (21) | 0 (0) | 0 (0) |
| ***S. sonnei (n = 115)*** | 110 (96) | 105 (91) | 92 (80) | 60 (52) | 112 (97) | 112 (97) | 101 (88) | 113 (98) |
| **EIEC (n = 31)** | 9 (29) | 5 (16) | 1 (3) | 2 (6) | 19 (61) | 11 (35) | 3 (10) | 4 (13) |
| **Other *E. coli (n =21)*** | 6 (29) | 10 (48) | 12 (57) | 2 (10) | 11 (52) | 9 (43) | 4 (19) | 4 (19) |
| **Unassigned** | 1 (0.4) | 1 (0.4) | 0 (0) | 0 (0) | 0 (0) | 2 (1) | 0 (0) | 0 (0) |

*Percentage correct identification of total isolates (n =269) is displayed.*
